# Modified viral-genetic mapping reveals local and global connectivity relationships of ventral tegmental area dopamine cells

**DOI:** 10.1101/2022.01.18.476718

**Authors:** Kevin T. Beier

**Affiliations:** Department of Physiology and Biophysics, Neurobiology and Behavior, Biomedical Engineering, Pharmaceutical Sciences, Center for the Neurobiology of Learning and Memory, University of California, Irvine, Irvine, CA, USA 92697

## Abstract

Dopamine cells in the ventral tegmental area (VTA^DA^) are critical for a variety of motivated behaviors. These cells receive synaptic inputs from over 100 anatomically-defined brain regions, which enables control from a distributed set of inputs across the brain. Extensive efforts have been made to map inputs to VTA cells based on neurochemical phenotype and output site. However, all of these studies have the same fundamental limitation that inputs local to the VTA cannot be properly assessed due to non-Cre-dependent uptake of EnvA-pseudotyped virus. Therefore, the quantitative contribution of local inputs to the VTA, including GABAergic, DAergic, and serotonergic, is not known. Here, we used a modified viral-genetic strategy that enables examination of both local as well as long-range inputs to VTA^DA^ cells. We found that nearly half of the total inputs to VTA^DA^ cells are located locally, revealing a substantial portion of inputs that have been missed by previous analyses. The majority of inhibition to VTA^DA^ cells arises from the substantia nigra pars reticulata, with large contributions from the VTA and the substantia nigra pars compacta. In addition to receiving inputs from VTA^GABA^ neurons, DA neurons are connected with other DA neurons within the VTA as well as the nearby retrorubal field. Lastly, we show that VTA^DA^ neurons receive inputs from distributed serotonergic neurons throughout the midbrain and hindbrain, with the majority arising from the dorsal raphe. Our study highlights the importance of using the appropriate combination of viral-genetic reagents to unmask the complexity of connectivity relationships to defined cells in the brain.

## INTRODUCTION

VTA^DA^ cells mediate a variety of motivated behaviors, including reward and aversion.(Björklund & Dunnett, 2007; Bromberg-Martin et al., 2010; Cohen et al., 2012; Lammel et al., 2012; Wise, 2004). Substantial effort has been made to map the brain regions and cell types that provide input to and receive projections from VTA^DA^ cells, a critical step towards understanding how VTA^DA^ cells effect behavioral consequences in response to a variety of stimuli. The recent advent of one-step rabies virus (RABV) has enabled the mapping of inputs onto defined cell types (Wickersham et al., 2007). This strategy was employed nearly a decade ago to map brain-wide inputs to DA cells in the VTA and the adjacent substantia nigra pars compacta (SNc) (Watabe-Uchida et al., 2012). However, DA neurons in the VTA and SNc are not homogenous, but rather are heterogenous in their molecular signatures, projection patterns, physiological properties, and behavioral functions (Kim et al., 2016; Lammel et al., 2008, 2011, 2012; Lerner et al., 2015). We therefore designed a method, Tracing the Relationship between Inputs and Outputs (TRIO) to map the input-output relationship of projection-defined VTA^DA^ cells (Beier et al., 2015; Lerner et al., 2015; Schwarz et al., 2015). TRIO revealed that different subtypes of VTA^DA^ cells received biased inputs, and that global input-output maps could be used to infer the behavioral contribution of particular input sites, such as the cortex (Beier et al., 2015).

While these and more recent studies have mapped global inputs to VTA^DA^ and SNc^DA^ cells (Beier et al., 2015; Faget et al., 2016; Lerner et al., 2015; Menegas et al., 2015; Watabe-Uchida et al., 2012) each study has the same limitation in that only inputs located at a distance from the injection site in the midbrain can be assessed. This is because the Cre-dependent TVA protein that facilitates EnvA-mediated infection is also expressed at low levels in non-Cre-expressing cells near the site of injection (Miyamichi et al., 2013). Therefore, RABV virions that are pseudotyped with the EnvA protein can infect both Cre-expressing starter cells as well as non-Cre-expressing cells nearby that express low levels of TVA through leaky, non-Cre-dependent expression. This leaky gene expression is due to incomplete suppression of transcription/translation of TVA. While this may not be an issue with fluorescent molecules or chemogenetic effectors such as DREADDs (Botterill et al., 2021), in cases where only small amounts of a gene product are required to exert function, such as TVA, the problem becomes magnified. Only a single functional unit – presumably three TVA molecules bound to an EnvA trimer (Alsteens et al., 2017) – is required to enable infection of EnvA-pseudotyped RABV. Injection of *AAV-CAG-FLEx^loxP^-TVA* and *AAV-CAG-FLEx^loxP^-RABV-G* 2 weeks prior to injection of EnvA-pseudotyped RABV can result in thousands of infection events near the injection site, even in Cre-animals that should not express TVA (Beier et al., 2015). Notably, these infections do not occur if AAVs are not injected, demonstrating that the background is TVA-mediated. Therefore, whether RABV-labeled neurons near the injection site were bona-fide inputs to starter cells or cells that were infected via the viral inoculum in previous VTA^DA^ mapping experiments cannot be distinguished.

One solution to this problem is to reduce the efficiency of TVA-mediated infection. A mutant version of TVA with a single point mutation (Glu^66^→Thr, or 66T) that exhibits only 10% of the efficiency of the wild-type TVA dramatically reduces local background (Miyamichi et al., 2013; Rong et al., 1998). Using this variant enables analysis of the quantitative contribution of local inputs to defined cell types, such as VTA^DA^ neurons. Here, we quantified the inputs to VTA^DA^ neurons, comparing long-distance inputs to local inputs. We used the *TC^66T^* variant in place of the wild type TVA, *TC^B^*, in our tracing studies and found that almost half of the inputs to VTA^DA^ cells are located near the VTA; many of these represent inputs that were missed or may have been mis-quantified in previous studies. We then performed quantitative analyses to identify the biases that local inputs have onto different sets of VTA^DA^ cells. We then examined the sources of local GABAergic input to VTA^DA^ cells, explored potential connections between DA cells in the ventral midbrain, and quantified the sources of serotonergic inputs to VTA^DA^ cells. Our analysis is the first of its kind to detail the quantitative contributions of inhibition and neuromodulatory influence from local cells in the midbrain and hindbrain onto VTA^DA^ cells, providing a comprehensive global picture of the main cell populations that influence and control VTA^DA^ cells.

## RESULTS

We performed RABV one-step mapping using *TC^66T^* in place of *TC^B^* (Miyamichi et al., 2013) (Figure 1A). We first injected a combination of Cre-dependent adeno-associated viruses (AAVs) encoding the RABV glycoprotein, *RABV-G*, and *TC^66T^* into the VTA of *DAT-Cre* mice that express the Cre recombinase in DA neurons. Two weeks later, we injected EnvA-pseudotyped, G-deleted, GFP-expressing RABV into the VTA. We then allowed 5 days for RABV spread to input cells before terminating the experiment.

**Figure 1:**
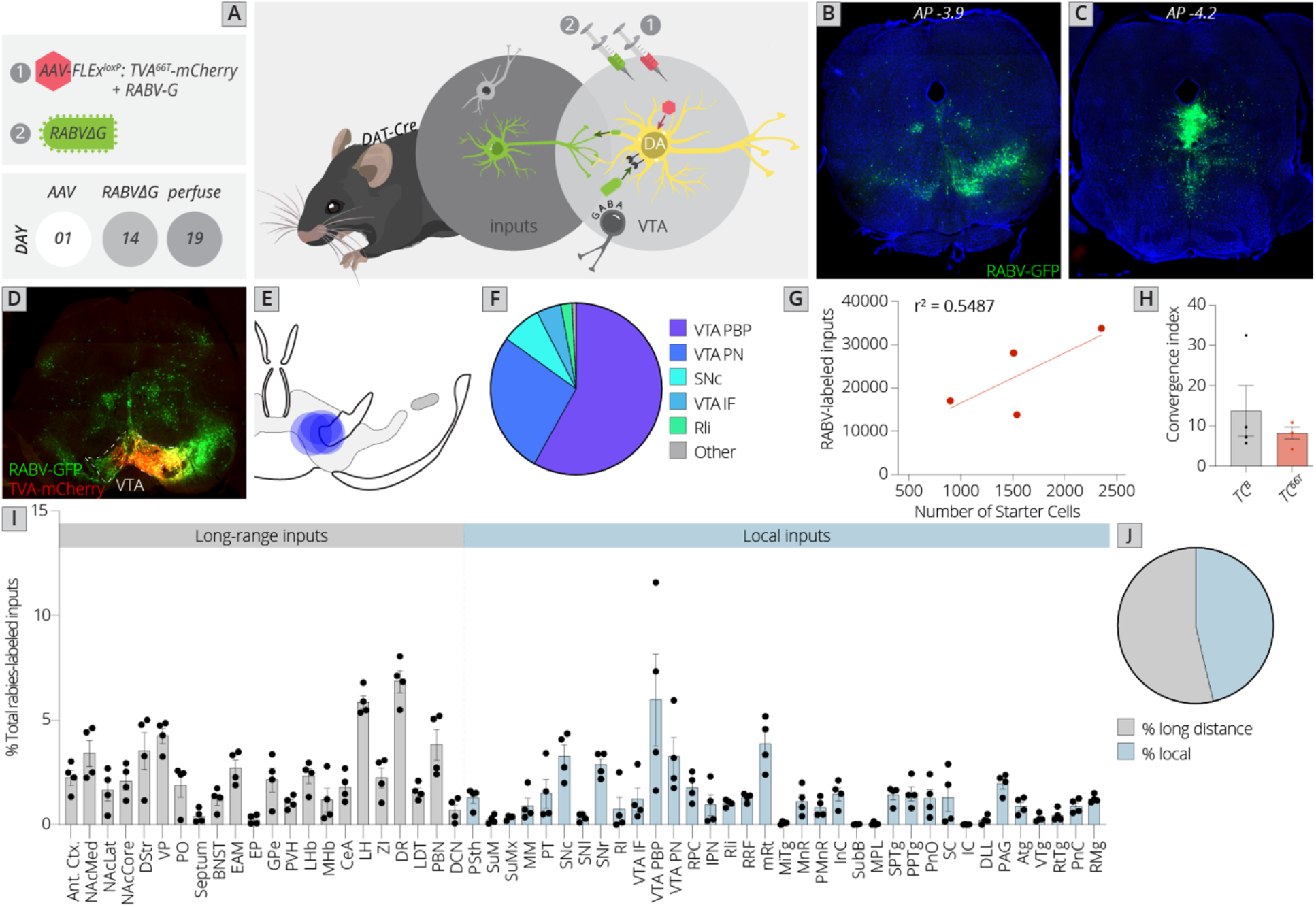
Use of a modified viral-genetic strategy to map local and global inputs to VTA^DA^ neurons. (A) Strategy for viral mapping. On day 1, a combination of AAVs expressing the mutated TVA protein fused to mCherry, *TC^66T^*, and RABV-G were injected into the VTA. Two weeks later, EnvA-pseudotyped RABV expressing GFP was injected into the VTA. Animals were sacrificed five days later. (B) Representative image of local virally-labeled inputs at anterior-posterior −3.9 mm from bregma. Scale, 1 mm. (C) Representative image of local virally-labeled inputs at anterior-posterior −4.2 mm from bregma. Scale, 1 mm. (D) Representative image of a midbrain section including starter cells in the VTA as well as local inputs. Scale, 1 mm. (E) Starter cell distributions for each of the four experiments. The center of each oval represents the center of mass of starter cells, and the horizontal and vertical radii of the oval represent one SD of starter cells in the medial-lateral and dorsal-ventral axes, respectively. (F) Fraction of starter cells located in each region in the ventral midbrain. 58 ± 6% were located in the PBP, 27 ± 6% in the PN, and 5 ± 2% in the IF nuclei of the VTA, 8 ± 3% in the SNc, 2 ± 1% in the Rli, and 1 ± 0.3% in all other regions. (G) Linear regression of RABV-labeled inputs vs. number of starter cells. (H) The convergence index, or ratio of inputs to starter cells, for RABV input mapping experiments using *TCB* as reported in Beier et al., *Cell* 2015, or *TC^66T^*. Only long-range inputs were considered to enable direct comparison between conditions. p = 0.42, 95% CI −21.25 to 10.24, unpaired t-test. (I) The percentage of each local and long-distance input relative to all RABV-labeled inputs is shown. (J) The fraction of inputs that were long-range inputs (54 ± 3%), as mapped by us in previous studies, and local inputs (46 ± 3%), is shown. All error bars in this figure and others throughout this manuscript represent ± 1 SEM.

Control experiments in non-Cre-expressing mice were performed to examine the local background of non-Cre-mediated, *TC^66T^*-facilitated infection. We previously observed using *TC^B^* that, on average, 3,183 cells were infected in a non-Cre-dependent, *TC^B^*-facilitated fashion per mouse (Beier et al., 2015). Using *TC^66T^*, we found an average of 2.67 cells per brain (Supplemental Figure 1). This background infection was indistinguishable from controls where no AAV was injected (Supplemental Figure 1). When experiments were performed in *DAT-Cre* mice, between 900 and 2400 starter cells were observed, and an average of 23,000 total input cells were labeled per experiment (Figure 1B-C). This resulted in a signal to noise ratio of 8,710 bona fide inputs: background infection. This is substantially higher than the signal: noise ratio obtained using *TC^B^* (approximately 13: see methods for quantification.)

**Supplemental Figure 1:**
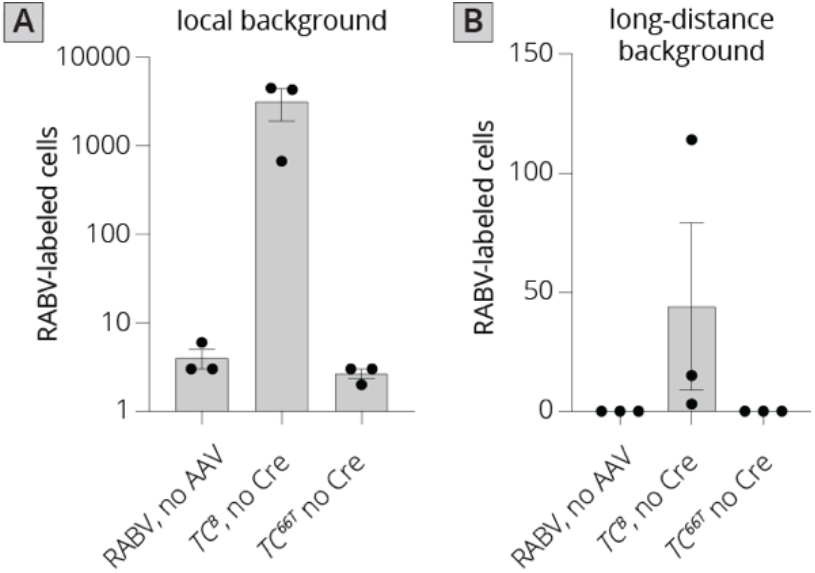
Quantification of local and long-distance input labeling using *TC^B^* and *TC^66T^*. (A) Local background in animals where AAVs were not injected in Cre+ mice, *TC^B^* was injected in Cre-mice, or *TC^66T^* was injected into Cre-mice. (B) Long-distance background in animals where AAVs were not injected in Cre+ mice, *TC^B^* was injected in Cre-mice, or *TC^66T^* was injected into Cre-mice.

We next examined the location of starter neurons within the ventral midbrain (Figure 1D-F). The majority of starter cells were located within the parabrachial pigmented nucleus (PBP: 58 ± 6%) and paranigral nucleus (PN: 27 ± 6%) of the VTA. The rest of the starter neurons were located within the medial aspect of the SNc (8 ± 3%), interfascicular nucleus of the VTA (IF: 5 ± 2%), and rostrolinear nucleus (Rli: 2 ± 1 %). We then examined the ratio of inputs to starter cells, also known as the convergence index, and compared it to tracing performed using *TC^B^*. The number of RABV-labeled inputs scaled with the number of starter cells, as expected (Figure 1G). The average convergence index using *TC^66T^* was approximately 8, which was not significantly different from tracing performed using *TC^B^* (Figure 1H), and also similar to the original study by Watabe-Uchida and colleagues using a non-mutated TVA, where the convergence index was approximately 7 (Watabe-Uchida et al., 2012). However, in our study nearly half (46%) of the inputs to VTA^DA^ cells arose from local regions that were not quantified in our previous analysis. Though Faget and colleagues excluded the VTA, SNc, SNr, red nucleus (RPC), and interpeduncular nucleus (IPN) likely for precisely the reason that concerned us, these regions combined to yield 20% of the total inputs to the VTA. This included about 10% of the total inputs within the VTA itself, 6% in the adjacent substantia nigra, and 4% in other nearby regions.

However, in addition to this 20% of inputs in regions immediately adjacent to the injection site, given that AAVs can exhibit substantial spread from the injection site we were concerned that non-Cre-dependent, EnvA-mediated infection extended beyond these regions to several other midbrain regions. To assess if this was indeed a problem, we compared our data to that published by Faget and colleagues for three separate groups of regions: 1) Those excluded in the previous study, 2) Midbrain/hindbrain regions excluded by us previously but not by the previous study, 3) long-range inputs included by both studies. Using dimensional reduction techniques to compare the overall labeling patterns within these three groups, we found that our data mixed with that from the Faget et al. study for long-range inputs (comparison 3) suggesting that the data were comparable. However, data from the two studies segregated for both the excluded (comparison 1) and questionable (comparison 2) brain regions (Supplemental Figure 2). These results support the likelihood that our method of using a modified, less efficient version of TVA is necessary to properly examine the local input landscape in the ventral midbrain.

### Local inputs to VTA^DA^ cells have differential associations with long-range inputs

Given the large fraction of total inputs to VTA^DA^ that are local, we wanted to explore the nature of these connections and by extension, how they may influence VTA^DA^ cells. Long-range inputs to DA neurons measured using RABV tracing have been published several times (Beier et al., 2015; Faget et al., 2016; Watabe-Uchida et al., 2012). More recently, DA neurons have been subdivided by output site, and the inputs to particular subpopulations compared to one another (Beier et al., 2015, 2019; Derdeyn et al., 2022; Lammel et al., 2012; Menegas et al., 2015).These analyses have enabled us to understand how input patterns relate to one another, and which inputs are biased onto which sets of VTA^DA^ cells. We can leverage these previous analyses to explore with which long-range inputs each set of local inputs associates, and by extension, which sets of DA cells may be predominately targeted by each set of inputs.

We first used Uniform Manifold Approximation and Projection (UMAP) to identify the clustering relationships of different input sites. We found that local input sites intermingled with long-distance inputs (Figure 2A). This is expected, and suggests that local inputs co-organize with long-distance inputs, rather than being a separate set of circuits. Given that we used only 4 brains for these UMAP analyses, which is a relatively small number for accurately capturing variance in the dataset, we wanted to test how robust our UMAP embedding was in capturing the relationships between brain regions. We recently performed a similar analysis on a 76 brain dataset that included long-distance RABV-labeled inputs to VTA cells based on projection, neurochemical phenotype, or a combination of these factors (Derdeyn et al., 2022). Since UMAP embeddings can be somewhat stochastic due to their reliance on initial seeding conditions, we also computed the distance between points relative to the maximum distance between any two points in each embedding, over 20 embeddings, then averaged across all embeddings for our four brain dataset, as done previously for the 76 brain dataset (Derdeyn et al., 2022). We found that the region associations were robust, with only three long-range input regions – the nucleus accumbens medial shell (NAcMed), nucleus accumbens core (NAcCore), and extended amygdala (EAM) associating with different groups of inputs across conditions (Figure 2B-C). While the results were not identical, these data suggest that our four brain dataset could be compared with reasonable confidence to our previous dataset.

**Figure 2:**
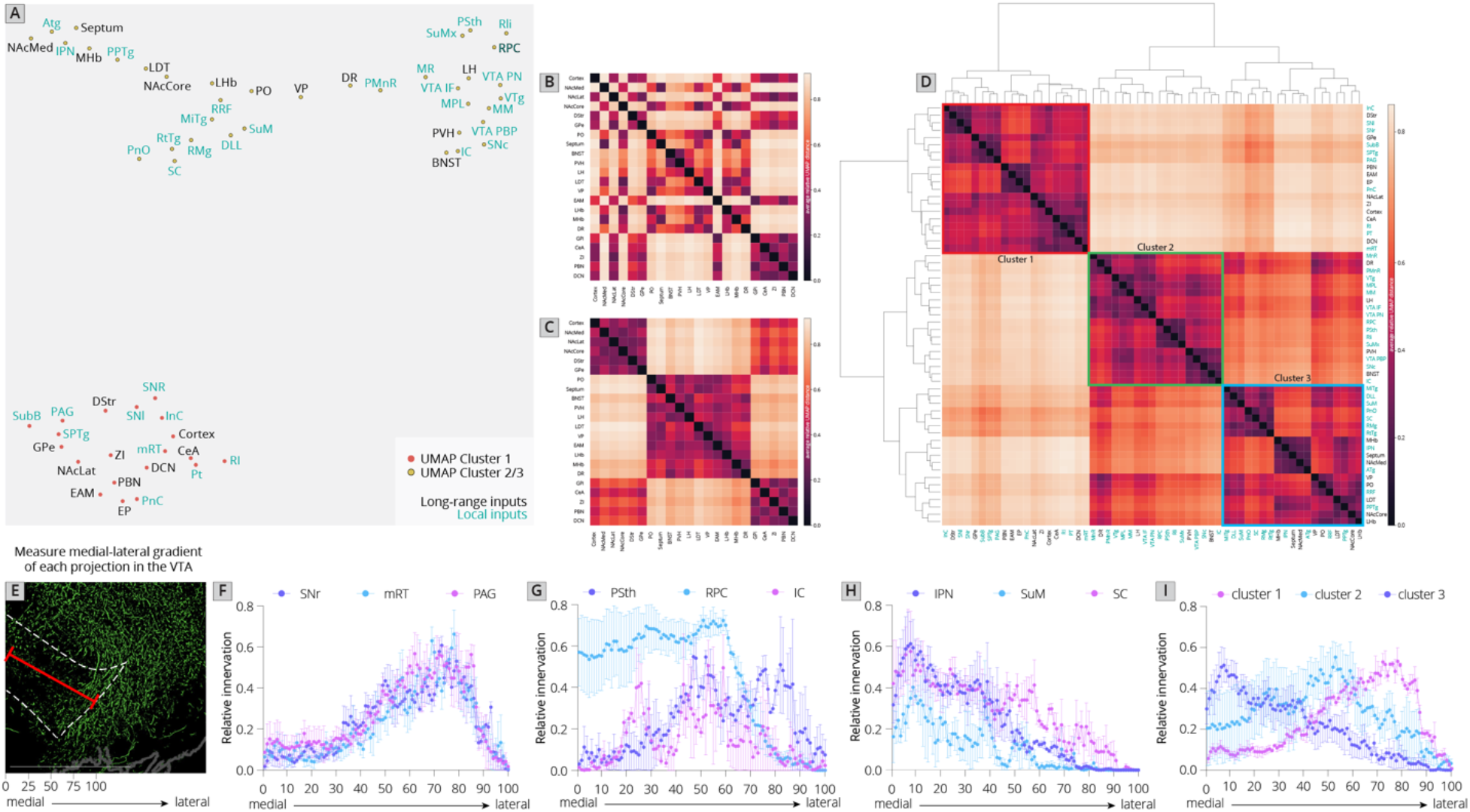
Association of local inputs to VTA^DA^ cells with long-range inputs. (A) Input regions are plotted in UMAP space, embedded with respect to z-scored counts across mouse brains. Clusters represent inputs with similar patterns of variation across the cohort. (B) Heatmap of pairwise distances (averaged across 20 UMAP embeddings) for the RABV input data, including only long-distance inputs to enable comparison to our 76-brain dataset, where only long-distance inputs were analyzed. (C) Heatmap of pairwise distances from an aggregated UMAP analysis of our 76 brain dataset, from Derdeyn et al. 2022, for purposes of comparison. (D) Heatmap of pairwise distances for the RABV input data, including both local and long-distance inputs. Regions are grouped according to hierarchical clusters. Clusters are highlighted to match the clusters in the UMAP plot. (E) Schematic of how relative innervation of the medial-lateral axis of the VTA was quantified. Scale, 1 mm. Images used for quantification of the data in panels E-I were obtained from the Allen Mouse Brain Connectivity Atlas. (F) Relative innervation across the medial-lateral axis for UMAP cluster 1. (G) Relative innervation across the medial-lateral axis for UMAP cluster 2. (H) Relative innervation across the medial-lateral axis for UMAP cluster 3. (I) Relative innervation across the medial-lateral axis for each UMAP cluster, representing the average of the three individual quantified input regions shown in F-H.

**Supplemental Figure 2:**
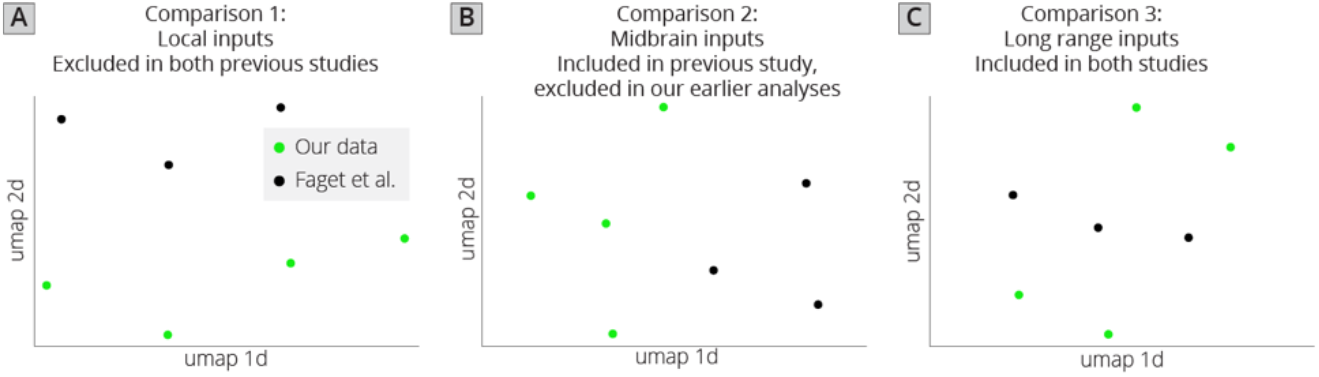
UMAP analysis of VTA^DA^ input tracing datasets. (A) UMAP embedding of brains using only inputs excluded in our previous studies as well as in Faget et al. These included the VTA, substantia nigra (SNc/SNr), IPN, RPC, and MHb. (B) UMAP embedding of local inputs included in Faget et al., but excluded from our previous studies. These inputs included the PSth, MM, SuM, MnR, PPTg, RMg, RRF, PT, InC, PAG, RI, VTg, colliculus (IC + SC), MiTg, SPTg, ATg, DLL, PnC, PnO, RtTg, and SubB. (C) UMAP embedding of long-range inputs from our study and Faget et al. Regions of the isocortex anterior to the corpus collosum were combined to make the anterior cortex. The EAM and DCN were excluded from the analysis as these were not contained in both datasets, and the MHb was excluded from this dataset because it was contained in the excluded dataset (panel A).

Similar to our 76 brain dataset, we observed three clusters of input regions (Figure 2D). The first is the set of brain regions that predominately project lateral or dorsal to the VTA (Derdeyn et al., 2022). The regions that project laterally include inputs from the basal ganglia such as the DStr, NAcLat, GPe, and cortex. Those that project dorsally include the entopeduncular nucleus (EP), zona incerta (ZI), and deep cerebellar nuclei (DCN). The local inputs that associate with this cluster include mostly those located dorsally and laterally to the VTA, including the periaqueductal gray (PAG), midbrain reticular nucleus (mRT), interstitial nucleus of Cajal (InC), subpeduncular tegmental nucleus (SPTg), subbrachial nucleus (SubB), SNr, and SNl. To test if the local inputs we identified indeed projected laterally to the VTA, we mapped out the relative innervation by the SNr, mRT, and PAG of the VTA across the medial-lateral gradient, as done previously for long-range inputs, using the Allen Mouse Brain Connectivity Atlas (Figure 2E) (Beier et al., 2019). We indeed observed that three selected inputs in cluster 1 – the SNr, mRT, and PAG - displayed a lateral bias in the VTA (Figure 2F, Supplemental Figure 3).

The second cluster of inputs includes mostly regions that project uniformly across the VTA, including the lateral hypothalamus (LH), paraventricular hypothalamus (PVH), and bed nucleus of the stria terminalis (BNST) (Derdeyn et al., 2022). The local inputs associating with this cluster include inputs from all of the subregions of the VTA itself (PBP, PN, IF), and other inputs located along the midline [median raphe (MnR), supramammillary decussation (SuMx), medial mammillary nucleus (MM)] as well as those located just laterally to the midline such as the red nucleus (RPC), ventral tegmental nucleus (VTg), inferior colliculus (IC), and parasubthalamic nucleus (PSth). While more varied in their individual projections, the PSth, RPC, and IC show a more consistent projection across the medial-lateral axis of the VTA (Figure 2G).

The final cluster of inputs mostly includes those that project medially within the VTA. This includes the MHb and lateral habenula (LHb), septum, preoptic nucleus (PO), and laterodorsal tegmentum (LDT). Local inputs associating with this group include midline structures such as the IPN, raphe magnus (RMg), supramammillary nucleus (SuM), structures near the midline such as the reticulotegmental nucleus (RtTg), anterior tegmental nucleus (ATg), superior colliculus (SC), slightly more lateral structures such as the retrorubal field (RRF) and pedunculopontine tegmental nucleus (PPTg), or inputs with lower counts such as the microcellular tegmental nucleus (MiTg) and the dorsal nucleus of the lateral lemniscus (DLL). The IPN, SuM, and SC all showed a medial bias in the VTA (Figure 2H). The difference in medial-lateral preference among the three clusters was especially clear when averaged across the three tested regions and plotted together (Figure 2I). These data in sum suggest a topographical organization of inputs to the VTA that applies to both global and local inputs, whereby inputs segregate based on their projection medially or laterally within the VTA, as we have found previously with long-range inputs (Beier et al., 2015, 2019; Derdeyn et al., 2022). Here, we extend these observations by including all inputs to the VTA, regardless of distance from injection site.

**Supplemental Figure 3:**
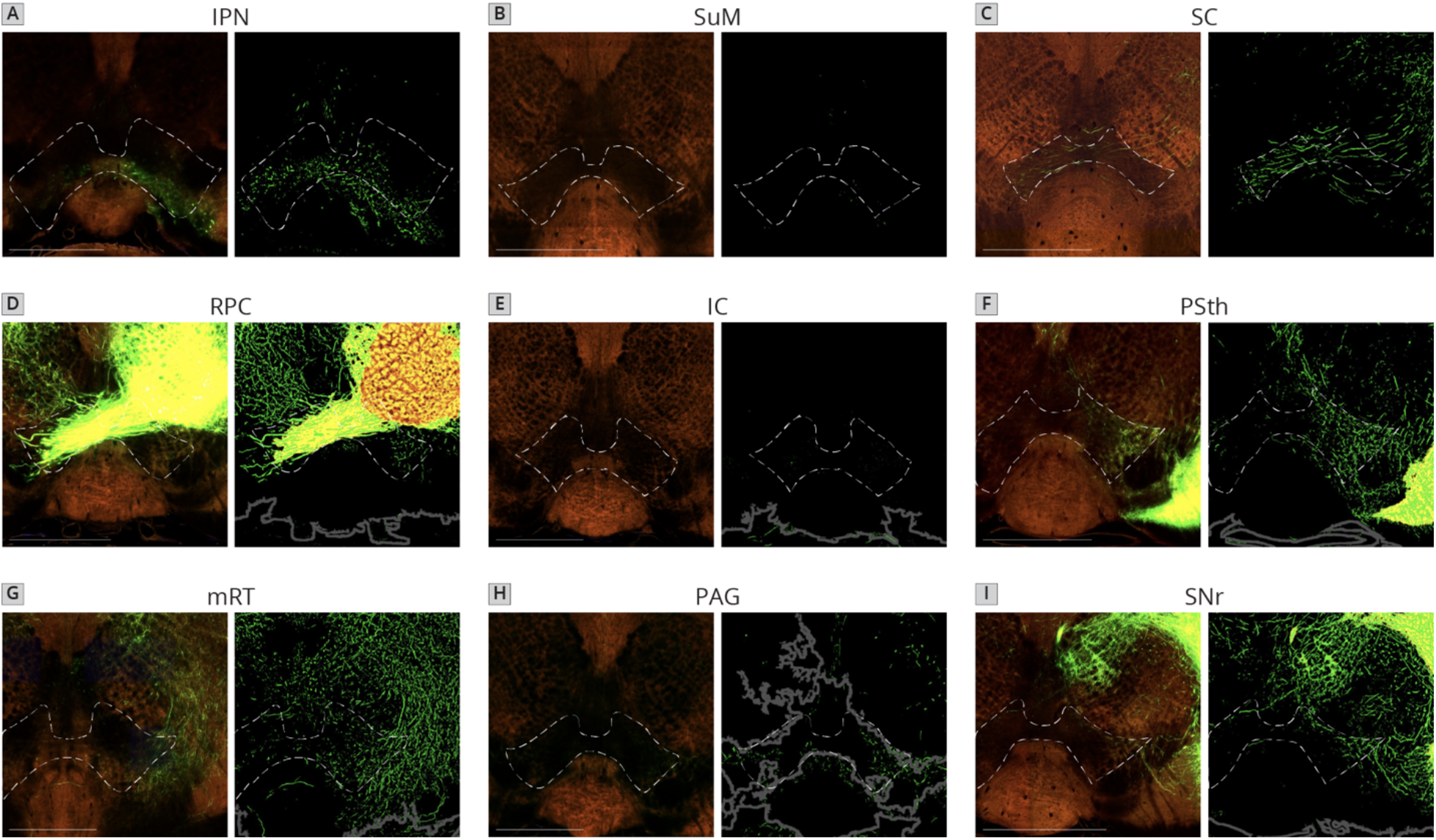
Sample images of projections in the VTA from each of the 9 quantified local inputs to VTA^DA^ cells. Images are from axons of labeled cells in the (A) IPN, (B) SuM, (C) SC, (D) RPC, (E) IC, (F) PSth, (G) mRT, (H) PAG, (I) SNr.

### Local and distributed GABAergic inhibition to VTA^DA^ cells

We next wanted to assess the location of local GABAergic inhibition to the VTA. GABA neurons functionally oppose DA neurons, providing a punishment and aversion signal (Bouarab et al., 2019; Tan et al., 2012). The majority of studies of inhibition to VTA^DA^ cells have focused on either GABA neurons within the VTA or on an anatomically poorly-defined structure referred to as the rostromedial tegmental nucleus (RMTg) (Jhou 2005; Jhou et al., 2009a; Jhou et al., 2009b; Kaufling et al., 2009). While extensive efforts have mapped the inputs, including inhibitory cell inputs to the VTA (Geisler et al., 2007; Geisler & Zahm, 2005; Phillipson, 1979; Sesack & Grace, 2010; Swanson, 2000; Zahm et al., 2011), these methods lacked the connectivity information afforded by RABV. We therefore performed our local mapping experiments in combination with fluorescent in situ hybridization against the GABAergic markers GAD1 and GAD2 in order to provide a quantitative map of inhibition onto VTA^DA^ cells (Figure 3A-B).

**Figure 3:**
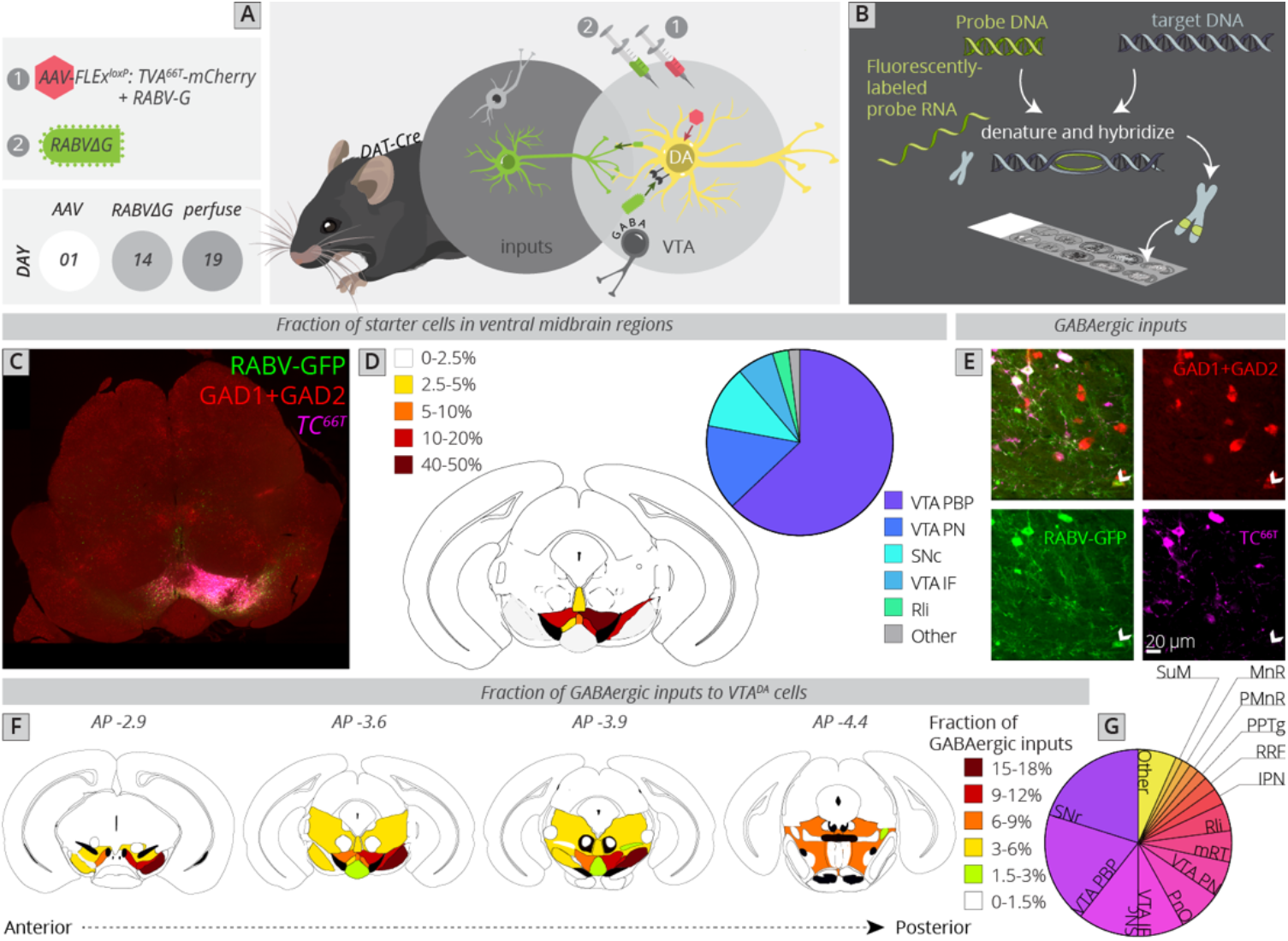
Local and distributed GABAergic inhibition to VTA^DA^ cells. (A) Strategy for viral mapping combined with fluorescent in-situ hybridization with probes for GAD1 and GAD2. (B) Sample histology section of the midbrain near the injection site. Green = RABV, red = FISH for GAD1/GAD2, magenta = *TC^66T^*. Scale, 1 mm. (C) Breakdown of starter cell location in different midbrain nuclei. 63 ± 1% were located within the PBP nucleus of the VTA, 15 ± 1% in the VTA PN, 11 ± 2% in the SNc, 6 ± 1% in the VTA IF, 3 ± 1% in the Rli, and 2 ± 0.3% in all other regions. (D) Example histology image of labeled cells, with the arrowhead pointing to a RABV+, GAD1/GAD2+, *TC^66T^* - cell. (E) Schematic heatmap of GABAergic input location throughout the ventral midbrain. Colors correspond to the total % of local GABAergic inputs located in that particular region. Data from each hemisphere is reported separately. The injection occurred in the right hemisphere. (F) Pie chart representing the total fraction of GABAergic inputs located in each region. Data from both hemispheres are combined. 20.1 ± 1.4% of all inhibitory inputs are from the SNr, 19.3 ± 0.7% from the VTA PBP, SNc 10.5 ± 2.4%, VTA IF 8.0 ± 2.6, VTA PN 6.2 ± 0.6, PnO 7.5 ± 2.4%, mRT 5.5 ± 1.3%, Rli 4.8 ± 0.5, IPN 2.8 ± 1.8, RRF 2.5 ± 1.1%, PPTg 1.8 ± 0.8%, PMnR 1.6 ± 0.3%, MnR 1.2 ± 1.2%, SuM 1.2 ± 0.2, and 6.6 ± 0.8 in all other regions.

In order to assess how this local input map would relate to our whole-brain local/global maps elucidated in Figure 1, we first compared the location of starter neurons in the VTA between experiments (Figure 3C-D). The majority of starter neurons were located within the PBP nucleus of the VTA (63 ± 1%), followed by VTA PN (15 ± 1%), SNc (11 ± 2%), VTA IF (6 ± 1%), and Rli (3 ± 1%). This was a similar distribution to our whole-brain mapping experiments (Figure 1F), enabling direct comparison of these datasets.

We counted all RABV-labeled neurons in regions local to the VTA, and assessed their co-staining with GAD1/GAD2 markers as well as *TC^66T^*. GABAergic input neurons were identified as those that were negative for *TC^66T^* and positive for GAD1/GAD2 (Figure 3E). We found that, surprisingly, the SNr is the single largest source of inhibition onto the cells that we targeted, representing 20.1 ± 1.4% of all inhibitory inputs (Figure 3F-G). Given that the VTA contains 84% of starter cells and SNc contained only 11% of starter cells, and most of these SNc starter cells were on the border of the lateral VTA/medial SNc, these SNr neurons almost certainly inhibit both SNc and VTA neurons. The majority of labeled SNr cells were located in the ventral-medial aspect of the SNr, near to the VTA. The second largest source of input was the PBP nucleus of the VTA (19.3 ± 0.7%), followed by the SNc (10.5 ± 2.4%). These three inputs represented 50% of the total inhibition of VTA^DA^ neurons. With the addition of the IF (8.0 ± 2.6) and PN (6.2 ± 0.6) nuclei of the VTA, 64% of inhibitory neurons are from regions within or immediately adjacent to the VTA.

In addition to inhibitory input from the above regions, VTA^DA^ neurons received inhibitory inputs from distributed sites across numerous regions posterior to the VTA. These included the pontine reticular nucleus, oral part (PnO: 7.5 ± 2.4%), mRT (5.5 ± 1.3%), Rli (4.8 ± 0.5), IPN (2.8 ± 1.8), RRF (2.5 ± 1.1%), PPTg (1.8 ± 0.8%), paramedian raphe nucleus (PMnR: 1.6 ± 0.3%), MnR (1.2 ± 1.2%), and SuM (1.2 ± 0.2). Several of these regions are thought to contribute to the RMTg, consistent with their known behavioral role in modulating VTA^DA^ cells. However, it is notable that at least quantitatively, they represent a minor fraction of total inhibitory inputs to VTA^DA^ cells.

### Midbrain DA neurons exhibit extensive interconnectivity

VTA^GABA^ neurons in total comprised about 10% of the total inputs to VTA^DA^ cells (Figure 1), and about 33% of the total local GABAergic input to the VTA^DA^ cells (Figure 3). However, in addition to the VTA^GABA^ input, there is evidence that VTA^DA^ cells also receive inputs from local glutamatergic (Dobi et al., 2010) as well as DA cells. Physiological evidence has shown that DA neurons release DA within the midbrain via somatodendritic release (Bayer & Pickel, 1990; Groves & Linder, 1983). Locally released DA then binds D2 autoreceptors expressed on DA cells, suppressing their activity. We wanted to test if DA neurons were connected by conventional means, and if so, what the topology of DA-DA neuron connectivity is in the ventral midbrain.

The challenge of assessing DA-DA connectivity is to distinguish cells that express TVA and thus could serve as starter neurons, and those that did not (and thus represent bona-fide inputs). We previously published the extent of spread of both *AAV_5_-CAG-FLEx^loxP^-TCB* and *AAV_8_-CAG-FLEx^loxP^-RABV-G* in the ventral midbrain given the exact injection coordinates used in this study, which provided a quantified radius of spread for each virus (Beier et al., 2015, 2019). We stained brain sections with both an anti-tyrosine hydroxylase (TH) antibody to label DA cells, as well as an mCherry antibody to delineate *TC^66T^*-expressing neurons (Figure 4A-E). We then assessed regions outside of the sphere of TVA spread that were either anterior in the VTA, posterior/medial or posterior/lateral in the VTA, as well as in the nearly RRF, and tested for how many neurons co-stained with TH. We found that about 40% of inputs in each region that did not express clear mCherry protein, even after antibody amplification, co-stained with TH (Figure 4F). That this number did not substantially differ regardless of whether the site was nearer to the injection site (anterior, posterior/medial) or further away (posterior/lateral, RRF) provides further evidence that RABV labeling was not due to direct TVA-mediated uptake, as in that case we would expect a higher percentage of TH+ cells in regions near the injection site. Our results suggest that in addition to a substantial GABAergic input from the VTA to local VTA^DA^ cells, VTA^DA^ neurons receive a large input from other DA cells that comprises approximately 4-5% of the total inputs to VTA^DA^ cells (11% of total inputs from the VTA/RRF x 40% that are TH+). These results suggest that there is indeed a substantial level of connectivity between DA neurons throughout the VTA and adjacent structures.

**Figure 4:**
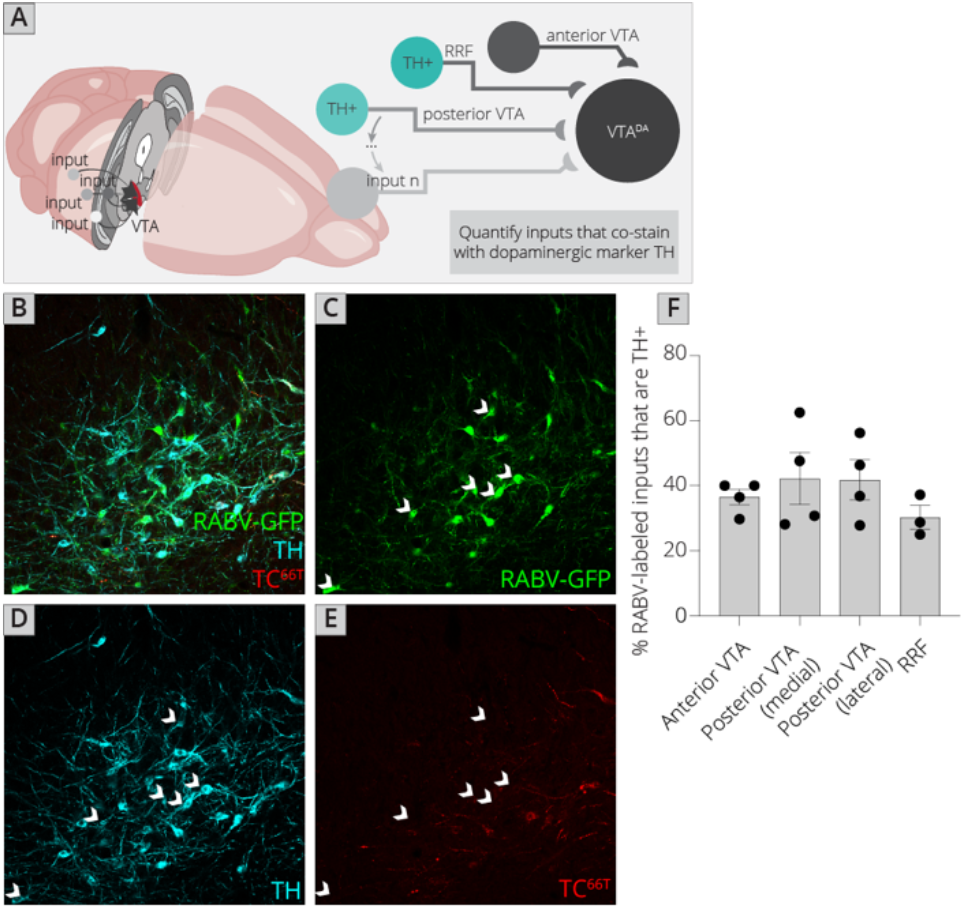
Connections between DA cells in the ventral midbrain. (A) Schematic of experiments for assessing potential DA-DA connections. Brain sections were stained for TH and mCherry and local inputs in the A8 (RRF) and A10 (VTA) regions were assessed for co-labeling with TH and mCherry. Cells that were GFP+, TH+, and mCherry-, were considered DAergic inputs. (B-E) Example histology image of labeled cells in the RRF. Green = GFP, cyan = TH, red = mCherry. Scale, 50 μm. (F) Quantification of TH+ inputs in four different midbrain regions.

### Sources of serotonergic inputs to VTA^DA^ cells

The brain’s neuromodulatory systems, including cells that release the monoamines DA, serotonin, or norepinephrine, are extensively interconnected (Hensler et al., 2013). In particular, serotonin (5-HT) plays a key role in the modulation of the DA system. A number of studies using classical methods such as small molecule and dye tracers have found that the VTA receives strong innervation from cells in the B7 region, also known as the dorsal raphe (DR) (Phillipson, 1979). However, the methods employed by these studies do not distinguish connected cells from passing fibers, or inputs connected to non-DA cells. Several studies including our own have demonstrated that serotonergic neurons in the DR do directly connect to both DA and GABA cells in the VTA (Beier et al., 2015; Liu et al., 2014; McDevitt et al., 2014; Qi et al., 2014; Wang et al., 2019). However, serotonergic input likely arises from other serotonergic nuclei besides the DR. Where the serotonergic neuron input is arising from, what the density of this innervation is, and the fraction of serotonergic innervation provided by each region is not known. Since several of the serotonergic cell-containing brain regions, including the pontine tegmentum (B9) and MnR (B8) regions are located nearby the VTA^DA^ cells, connectivity could not be discerned using viral mapping studies that employ the standard *FLEx^loxP^/DIO* TVA to mark starter cells. Therefore, our approach is uniquely suited to address this question.

We mapped inputs to VTA^DA^ neurons, as before, and co-stained neurons in the midbrain and hindbrain with an antibody to tryptophan hydroxylase 2 (Tph2), which marks serotonergic neurons (Figure 5A-B). We examined both hemispheres independently, except for the RMg (B3), DR (B7), and MnR (B8), as these cell populations are located along the midline. We observed that approximately 30% of cells in each region co-stained with Tph2, with a high of 37% and a low of 20% (Figure 5C). This number did not vary substantially, regardless of the region or hemisphere, suggesting that the fraction of cells providing direct connections onto VTA^DA^ neurons that are serotonergic within each region is relatively constant across all serotonergic nuclei. We also want to assess the fraction of total input provided by each set of serotonergic cells. We therefore normalized the number of serotonergic inputs in each region per 10,000 RABV-labeled input cells. This analysis showed that the vast majority of serotonergic inputs to the VTA indeed arise from the DR (376 ± 24 per 10,000 inputs; Figure 5D). The regions providing the next largest quantitative serotonergic inputs are the mRT (B6; 50 ± 11 inputs per 10,000), MnR (37 ± 4 inputs per 10,000), RMg (30 ± 6 inputs per 10,000), and pons (B5; 26 ± 4 inputs per 10,000). Therefore, while the majority of serotonergic inputs do arise from the DR, VTA^DA^ cells also receive distributed serotonergic input from several other midbrain and hindbrain nuclei that together represent approximately 1.5% of total inputs to VTA^DA^ cells.

**Figure 5:**
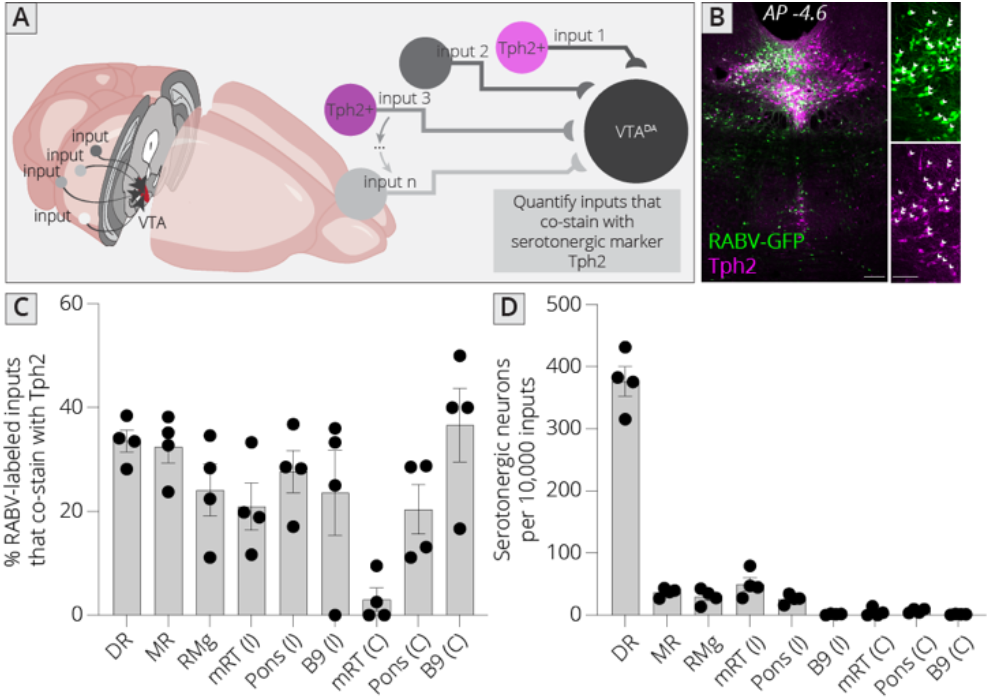
Identifying location of serotonergic inputs to VTA^DA^ cells. (A) Schematic of experiments for assessing potential serotonergic inputs. Brain sections were stained for Tph2, and inputs in the B3, B5, B6, B8, and B9 regions were assessed for co-labeling with Tph2. Cells that were GFP+ and Tph2+ were considered serotonergic inputs. (B) Example histology image of labeled cells in the midbrain. Green = GFP, magenta = Tph2. Scale, 50 μm. (C) Percentage of RABV-labeled inputs in each region that co-stained with Tph2. (D) The number of serotonergic inputs in each quantified region, normalized to 10,000 RABV-labeled inputs.

## DISCUSSION

In this study, we provide a comprehensive local and global input map to VTA^DA^ neurons. We focused on local inputs to the VTA that were inaccessible using previous approaches, and quantified where GABAergic, DAergic, and serotonergic inputs arose in the midbrain and hindbrain. Our study provides the first comprehensive map of these connections to VTA^DA^ neurons and thus provides a valuable resource for future studies of the VTA as well as a template for mapping both local and global inputs to other structures in the brain.

Inputs to DA cells in the VTA have been previously mapped (Beier et al., 2015; Faget et al., 2016; Watabe-Uchida et al., 2012). However, each of these studies reported a limited number of input regions, as the issue of local background was likely recognized in each study. The report from Watabe-Uchida and colleagues included a much reduced local input labeling relative to this study, perhaps due to the limited reporting of input regions. Most importantly, nearly every study to date, including the three cited above, used a wild-type version of the TVA protein that would likely yield hundreds to thousands of non-Cre-dependent, TVA-facilitated infections in the midbrain near the injection site. Our approach using *TC^66T^* to reduce non-Cre-mediated infections near the site of injection enabled high-resolution, low-noise analysis of connectivity to VTA^DA^ cells. The fact that nearly 50% of inputs are from local brain sites highlights the importance of local connectivity within the midbrain. That many of the brain regions are much less studied than their forebrain counterparts represents an opportunity to further study inputs from the midbrain and hindbrain that provide quantitatively substantial inputs to VTA^DA^ neurons yet whose functions remain largely unknown.

### Relationships with different long-range input sites

We previously mapped inputs to VTA neurons based on output site and neurochemical phenotype, amassing a 76 brain dataset that enabled us to dissect the sources that influence patterns of inputs to different cells in the VTA (Beier et al., 2015, 2019; Derdeyn et al., 2022). In this study, we wanted to map the local inputs onto these scaffolds to observe if local inputs have biases towards specific VTA^DA^ subpopulations. We demonstrated here using a subset of local input regions that the three clusters of input regions arising from our UMAP analysis were related to spatial projection to the VTA. We used output tracing experiments from three representative local input regions from the Allen Mouse Brain Connectivity Atlas, as the number we could select was quite limited. This was due to the fact that most of the local input brain regions are small, and thus without defined Cre cell populations to target each population, more often than not the desired brain region represented only a small fraction of the total cells expressing GFP. Even so, the brain regions that we identified that did contain at least 2 injections relatively specifically targeting the desired regions reinforced our interpretation that the UMAP clusters represent different medial-lateral projection patterns in the VTA.

We had previously linked biases in long-range inputs to different VTA cell populations, and showed that these biases were related to stereotyped projection archetypes in the VTA (Derdeyn et al., 2022). Namely, inputs that project laterally to the VTA predominantly innervate VTA^DA^→NAcLat cells, those that uniformly innervate the VTA are biased onto VTA^DA^→Amygdala cells, those projecting ventromedially preferentially target VTA^DA^→NAcMed cells, and those that project ventromedially target VTA^DA^→medial prefrontal cortex (mPFC) cells (Derdeyn et al., 2022). Though the four brains used for this analysis were many fewer than the 76 brain dataset used previously, we were able to recapitulate the main associations between brain regions, supporting the validity of our interpretations. This thus suggests that local inputs to cluster 1 (SNr, SNl, mRT, SPTg, SubB, PnC, RI, PT, InC) principally target VTA^DA^→NAcLat cells, those in cluster 2 (PSth, IC, RPC, VTA PBP, PN, IF, MnR, Rli, SNc, PMnR, VTg, MPL, MM, SuMx) predominately innervate VTA^DA^→Amygdala neurons, and those in cluster 3 (IPN, SuM, SC, PnO, ATg, RMg, PPTg, RRF, RtTg, MiTg, DLL) are biased onto VTA^DA^→NAcMed or VTA^DA^→mPFC cells. One important limitation is that our analyses here are limited to the medial-lateral axis in the VTA. We previously showed that projections demonstrate both a medial-lateral and dorsal-ventral gradient in the VTA (Derdeyn et al., 2022). We chose to focus here on the medial-lateral axis as that is the principal axis of variation for long-range inputs in the VTA (Derdeyn et al., 2022). Our analyses have generally not had sufficient anterior-posterior resolution since the location of starter cells, which was critical to the original observation of spatial gradients in inputs to the VTA, was obtained using serial 60 μm coronal sections in the VTA (Beier et al., 2015, 2019). Therefore, the local inputs mapped here may have additional gradients along the dorsal-ventral or anterior-posterior axes that were not captured in our analysis.

### DA-DA cell interconnectivity

In addition to local GABAergic inputs, we also found a large number of DA-DA cell connections in the VTA. DA neurons are known to signal to one another through release of DA, which is thought to signal through volume transmission (Groves & Linder, 1983; Hajdu et al., 1973; Wilson et al., 1977). It is therefore of interest to consider, given that the dominant hypothesis is that RABV is transmitted via synapses, how this DA-DA neuron transmission may occur. In a study mapping connections to direct and indirect pathway neurons in the dorsal striatum, RABV was also observed to transmit to DA cells (Wall et al., 2013). However, this mode of transmission occurs approximately 10-fold less efficiently than when RABV was injected directly into the dorsal striatum. However, it is not certain that inter-midbrain DA transmission occurs via classic volume transmission and not focal release/signaling of DA. For example, DA levels appear to rise to ≥ 10 uM for brief periods of ≤ 100 ms (Ford et al., 2009). This rapid on/off kinetics of DA signaling is inconsistent with DA signaling at a distance (Beckstead et al., 2004). Furthermore, spontaneous exocytotic, GPCR-mediated signaling was observed in SNc^DA^ neurons (Gantz et al., 2013). This evidence indicates that, like traditional ligand-gated transmission, GPCR-mediated transmission can occur in a point to point fashion, similar to a classic synaptic mechanism. More recent evidence using the DA sensor dLight and a photoactivatable D2 receptor ligand also supports the hypothesis that DA release occurs from highly specific sites (Condon et al., 2021). From our experiments we cannot infer the nature of DA release or contacts between DA cells. However, our results also support the possibility of specific contacts between DA cells through which RABV transmission can occur. We estimate that approximately 4-5% of total RABV-labeled inputs are from DA cells. If this number is also influenced by a ten-fold reduction in efficiency of transmission, as estimated from Wall and colleagues, this would indicate that the presence of DA terminals, and thus the potential for DA influence of other VTA^DA^ cells is extraordinarily high.

This interconnectivity of DA cells throughout the ventral midbrain also has implications for tract tracing studies. Several studies, including the earliest input-output viral-genetic mapping study of the VTA defined output cells by injection of RABV into a projection site, and enabled spread of RABV from VTA cells via injection of an AAV expressing RABV-G into the ventral midbrain of *DAT-Cre* mice (Lammel et al., 2012). According to this strategy, any DA cell expressing RABV-G could serve as a starter cell. Thus, for example when RABV was injected into NAcLat, if VTA^DA^→NAcLat cells receive input from VTA^DA^→mPFC cells, then RABV could spread to VTA^DA^→mPFC cells and subsequently, their inputs. This interconnectivity thus has the potential to degrade pathway-specific connectivity. Our TRIO strategy was designed to avoid this potential degradation by injection of a third virus, which we chose to be the canine adenovirus (CAV-2) (Beier et al., 2015; Lerner et al., 2015; Schwarz et al., 2015) and later AAV_retro_ or herpes simplex virus (Ren et al., 2018). We employed a dual recombinase AND-gate where cell types were defined by Cre expression and outputs defined by expression of Cre-dependent Flp recombinase, delivered via CAV-2. Our results from this study suggest that the AND-gate TRIO strategy is required for maintaining pathway specificity in input-output analysis.

### Distributed serotonergic inputs to VTA^DA^ cells

In addition to DA, serotonin plays a key role in the modulation of VTA^DA^ cells. Multiple types of serotonin receptors are expressed in the VTA, including 5-HT1B, 5-HT2A, and 5-HT2C (Bubar & Cunningham, 2007; Doherty & Pickel, 2000; Pazos & Palacios, 1985; Waeber et al., 1989). Application of serotonin to the VTA depolarizes dopamine neurons, likely through 5-HT2A receptors (Paolucci et al., 2003; Pessia et al., 1994). Studies using classical methods including horseradish peroxidase and radiolabeling found that the VTA receives inputs from brain regions that include serotonergic neurons, including most prominently the DR (Azmitia & Segal, 1978; Hervé et al., 1987; Parent et al., 1981; Phillipson, 1979). Our results using one-step RABV reinforce these previous findings. However, we also found that DA neurons receive direct innervation from serotonergic neurons located in several other regions, including the B3, B5, B6, B8, and B9 regions. The sum of these inputs equates to approximately 1.5% of total inputs to VTA^DA^ cells. While this may seem insubstantial, serotonergic inputs from the DR only represent about 3.7% of total inputs to VTA^DA^ cells, yet these provide a powerful input to VTA^DA^ cells that influences reward behavior by triggering DA release in the NAc (Liu et al., 2014; McDevitt et al., 2014; Qi et al., 2014; Wang et al., 2019). It is also possible that this underrepresents the total serotonergic influence on VTA^DA^ cells, as serotonin signaling, like DA, is thought to occur in part by volume transmission, therefore likely reducing the efficiency of RABV labeling. Notably, many serotonergic neurons in the DR also co-release glutamate, which contributes to the rewarding phenotype of stimulation. It is possible that serotonergic neurons in other brain sites may co-release other neurotransmitters as well, which may work in concert with serotonin to modulate the activity and function of VTA^DA^ cells.

### Practical concerns about local labeling without *TC^66T^*

The use of *TC^66T^* enables analysis of inputs near to the site of injection. This approach is required due to high levels of non-Cre-mediated, TVA-facilitated infection of EnvA-pseudotyped RABV. This highlights the issue that investigators regularly analyze brain regions near the injection site without performing the proper controls. The issue is not with the particular AAV vectors used for RABV mapping, but are representative of problems inherent in all standard AAV vectors. While FLEx^loxP^ and DIO cassettes are typically sufficient to silence expression of fluorescent reporter genes and functional effectors such as DREADDs or optogenetic proteins, low levels of non-Cre-mediated expression are a problem when only a small amount of gene product is required, such as in the case of recombinase proteins or viral receptors (Botterill et al., 2021; Miyamichi et al., 2013). In this case, even a low level of expression is sufficient to degrade specificity. Note that in control experiments where Cre-dependent TVA and RABV-G were introduced in non-Cre-expressing mice, we observed many fewer RABV-labeled cells located at a distance from the injection site. These results suggest that the leak of TVA was sufficient to mediate a substantial amount of infection of RABV locally, as well as some inputs located at a distance from the injection site. Given that we did not observe long-distance inputs labeled when AAV was not injected into the ventral midbrain, it is likely that labeled cells located at a distance from the midbrain were due to retrograde uptake of the TVA-expressing virus, and non-Cre-mediated expression of TVA from these inputs.

Given that nearly all RABV mapping experiments use wild-type TVA and many analyze virally-labeled cells near the injection site, it is important that the issue of non-recombinase-mediated, TVA-facilitated infection of EnvA-pseudotyped RABV is recognized by the community. As with other techniques, application of proper controls should be required for proper interpretation of RABV mapping experiments. We hope that our study serves as an exemplar for the application of such controls and the kinds of questions that can be investigated using the appropriate combination of viral vectors, as we used here to elucidate the nature of inhibitory and neuromodulatory inputs to DA cells in the VTA.

## Acknowledgements

We would like to thank Pieter Derdeyn for assistance with UMAP analysis, and May Hui for assistance with figure production. This work was funded by NIH DP2-AG067666, R00-D041445, R01-DA054374, TRDRP T31KT1437, and T31IP1426, One Mind OM-5596678, Alzheimer’s Association AARG-NTF-20-685694, New Vision Research CCAD2020-002, and ADPA APDA-5589562.

## MATERIALS AND METHODS

### Mice

*DAT-Cre* mice were obtained from the Jackson Laboratories (Backman et al., 2006). Mice were housed on a 12 hr light/dark cycle with food and water ad libitum. Viral vectors were prepared as previously described (Schwarz et al., 2015). All procedures followed animal care and biosafety guidelines approved by the University of California, Irvine’s Administrative Panel on Laboratory Animal Care and Administrative Panel of Biosafety. Both males and females were used in all experiments in approximately equal proportions.

### Viral tracing

One-step RABV input mapping was performed as previously described, with the substitution of *AAV_5_-CAG-FLEx^loxP^-TC^66T^* in place of *AAV_5_-CAG-FLEx^loxP^-TC^B^*(Beier et al., 2015). Briefly, 100 nL of a 1:1 volume mixture of *AAV_5_-CAG-FLEx^loxP^-TC^66T^* and *AAV_8_-CAG-FLEx^loxP^-RABV-G* was injected into the VTA of 6-week-old mice. Two weeks later, EnvA-pseudotyped, GFP-expressing RABV was injected into the VTA. Mice were allowed to recover for 5 days to enable viral spread, after which time experiments were terminated.

The titers of viruses, based on quantitative PCR analysis, were as follows: *AAV_5_-CAG-FLEx^loxP^-TC^66T^*, 2.4 × 10^13^ genome copies (gc)/mL *AAV_8_-CAG-FLEx^loxP^-RABV-G*, 1.0 × 10^12^ gc/mL.

The titer of EnvA-pseudotyped RABV was estimated to be 5.0 × 10^8^ colony forming units (cfu)/mL based on serial dilutions of the virus stock followed by infection of the 293-TVA800 cell line.

### Quantification of signal to noise ratio of RABV input mapping

In order to quantify the signal to noise ratio in *TC^66T^* vs *TC^B^*-mediated tracing, we considered 1) the number of neurons labeled local to the injection site in Cre-animals, which was considered the local background, 2) both the long-range and local inputs. We assumed that local inputs would comprise 46% of the total inputs for both *TC^B^* and *TC^66T^*-mediated tracing; therefore we multiplied the total inputs for *TC^B^* experiments by a constant factor (1.87) that was equal to the total inputs divided by the long-range inputs from *TC^66T^* tracing experiments. This resulted in an average total for *TC^B^*-mediated tracing of 40,955 inputs.

For *TC^B^* mapping experiments, we counted on average 3,183 local background cells and estimated a total of 40,955 RABV-labeled inputs. Therefore, this resulted in a signal/noise ratio of 40,955/3,183 = 12.9. For *TC^66T^* mapping experiments, we counted on average 2.67 local background cells and an average of 23,170 inputs. This yields a signal: noise ratio of 8,710.

### Immunohistochemistry

Animals were transcardially perfused with phosphate buffered saline (PBS) followed by 4% formaldehyde. Brains were dissected, post-fixed in 4% formaldehyde for 12–24 hours, and placed in 30% sucrose for 24–48 hours. They were then embedded in Tissue Freezing media and stored in a –80°C freezer until sectioning. For RABV tracing analysis, consecutive 60-μm coronal sections were collected onto Superfrost Plus slides and stained for NeuroTrace Blue (NTB, Invitrogen). For NTB staining, slides were washed 1×5 min in PBS, 2×10 min in PBS with 0.3% Triton X-100 (PBST), incubated for 2–3 hours at RT in (1:500) NTB in PBST, washed 1×20 min with PBST and 1×5min with PBS. Sections were additionally stained with DAPI (1:10,000 of 5 mg/mL, Sigma-Aldrich), which was included in the last PBST wash of NTB staining. Whole slides were then imaged with a 4x objective using an IX83 Olympus microscope.

For starter cell identification, sections were unmounted after slide scanning, blocked in PBST and 10% NGS for 2–3 hours at room temperature, and incubated in rat anti-mCherry antibody (1:2000, Life Sciences) and rabbit anti-TH antibody (1:1000, Millipore) at 4°C for four nights. After primary antibody staining, sections were washed 3×10 min in PBST, and secondary antibodies (donkey anti-rat Cy3 and donkey anti-rabbit 647, Jackson ImmunoResearch) were applied for two nights at 4°C, followed by 3×10min washes in PBST and remounting. Confocal z-stacks were acquired using a 20x objective on a Zeiss LSM 780 confocal microscope.

### Fluorescent In Situ Hybridization (FISH)

We performed FISH experiments as previously described (Weissbourd et al., 2014). To make ISH probes, DNA fragments of 400-1000 bp containing the coding or untranslated region sequences were amplified by PCR from mouse whole brain cDNA (Zyagen) and subcloned into pCR-BluntII-topo vector (Life Technologies, cat# K2800-20). The T3 RNA polymerase recognition site (AATTAACCCTCACTAAAGGG) was added to the 3’-end of the PCR product. Plasmids were then amplified, the insert removed via EcoRI (New England Biolabs, cat #R0101L) digest, and purified using a PCR purification kit (QIAGEN, cat #28104). 500-1000 ng of the DNA fragment was then used for in vitro transcription by using DIG RNA labeling mix (cat #11277073910) and T3 RNA polymerase (cat #11031163001) according to the manufacturer’s instruction (Roche Applied Science). After DNase I (Roche Applied Science, cat #04716728001) treatment for 30 min at 37°C, the RNA probe was purified by ProbeQuant G-50 Columns (GE Healthcare, cat# 28-9034-08) according to the manufacturer’s instructions. 60-μm consecutive sections were collected onto Superfrost slides (no-coating, Fisher Scientific, cat #22-034-980), dried, and stored at −80°C until use. Specific slides were then thawed and viewed on an Olympus compound fluorescence microscope, and the sections containing regions of interest were recorded. Those sections were then floated off using PBS into wells of a 24-well plate. The sections were fixed for 15 min in 4% formaldehyde in PBS at room temperature, rinsed with PBS, and incubated with 7 μg/ml Proteinase K (Life Technologies, cat #25530-049) in 10 mM Tris-Cl, pH 7.4, 1 mM EDTA for 10 min at 37°C. After fixing again with 4% formaldehyde in PBS for 10 min and rinsing with PBS, the sections were incubated with 0.25% acetic anhydride in 0.1 M triethanolamine, pH 8.0, for 15 min and washed with PBS. Probes were diluted (~1:1000) with the hybridization buffer (50% formamide, 10mM Tris-Cl pH 8.0, 200μg/ml tRNA, 10% Dextran Sulfate, 1x Denhalt’s solution, 600mM NaCl, 0.25% SDS), mixed well, preheated at 85°C for 5 min, and applied to each well (300-500 μl/well). After 16-20h of incubation at 50°C, the sections were washed, first with 2× SSC-50% formamide, then with 2× SSC, and finally with 0.2× SSC twice for 20 min at 65°C. After blocking for 1-2h with the 1% blocking reagent (Roche Applied Science, cat# 10057177103), sections were incubated with alkaline phosphatase-conjugated anti-DIG antibody (1:1000, Roche Applied Science, cat# 1093274) and chicken anti-GFP antibodies (1:500; Aves Labs, cat# GFP-1020) overnight at 4°C. After washing with Roche Wash Buffer (cat# 11585762001) three times for 15 min followed by rinsing with the detection buffer (100mM Tris-Cl pH8.0, 100mM NaCl, 10mM MgCl2), probe-positive cells were detected by Fast Red TR/Naphthol AS-MX Tablets (Sigma-Aldrich, cat# F4523). After washing with Roche Wash Buffer three times for 10 min, sections were incubated with FITC-conjugated donkey anti-chicken antibodies (1:200; Jackson ImmunoResearch) for an additional 1-2h, and washed with PBS three times for 10 min. Finally, the sections were treated with PBS containing 4’,6-diamidino-2-phenylindole dihydrochloride (DAPI; Sigma-Aldrich, cat# D8417) for 20 min and mounted with cover glass using Fluorogel (Electron Microscopy Sciences, Cat#17985-10). Sections were imaged by confocal microscopy (Zeiss 780). Images were processed in ImageJ. We used the Cell Counter plugin in FIJI to quantify overlap.

#### Dimensional reduction of RABV input data

Uniform Manifold Approximation and Projection (UMAP) was used as a non-linear dimensional reduction technique on input data. UMAP is optimized for finding local and global structures in high dimensional data. Analyses were performed using the official UMAP library (McInnes et al., 2018). The fractional counts data were z-scored to compare variation in output and input sites across regions with different magnitudes of counts. Z-scored data were dimensionally reduced with UMAP to find clusters of input sites with similar patterns of variation. UMAP parameters were tuned manually to optimize stability of clusters.

#### Analysis of fluorescent axonal labeling from Allen Mouse Brain Connectivity Atlas

Analysis was performed as previously described (Beier et al., 2019). Data from three separate injections for each local VTA^DA^ input site were analyzed. Experiments had to show GFP labeling in the VTA. For each experiment, three images of the VTA, each spaced approximately two sections apart, were captured at screen resolution. In ImageJ, using the line tool at a thickness of 100, a line was drawn from the midline to the end of the medial lemniscus running ventro-lateral through the VTA, and the gray value (a metric of axon coverage) was obtained as a function of distance from the midline. To normalize these values, data were run through custom MATLAB code to segment data into 100 bins and normalized to the maximum intensity value for that image. The three normalized values were then averaged for each brain, and the data from three separate brains then were averaged into one condition and plotted as a percentage across the medial-lateral axis.

#### Brain Region Abbreviation List

Abbreviations for brain regions made throughout the paper are listed below, in alphabetical order:

Long-range inputs:

Ant. Ctx./cortex: anterior cortex
BNST: bed nucleus of the stria terminalis
CeA: central amygdala
DCN: deep cerebellar nucleus
DR: dorsal raphe
DStr: dorsal striatum
EAM: extended amygdala
EP: entopeduncular nucleus (GPi)
GP: globus pallidus (GPe)
LDT: laterodorsal tegmentum
LH: lateral hypothalamus
LHb: lateral habenula
MHb: medial habenula
NAcCore: nucleus accumbens, core
NAcMed: nucleus accumbens, medial shell
NAcLat: nucleus accumbens, lateral shell
PBN: parabrachial nucleus
PO: pre-optic area
PVH: paraventricular hypothalamus
VP: ventral pallidum
VTA: ventral tegmental area
ZI: zona incerta

Local inputs:

ATg: Anterior Tegmental Nucleus
DLL: Dorsal Nucleus of the Lateral Lemniscus
IC: Inferior Colliculus
InC: Interstitial Nucleus of Cajal
IPN: Interpeduncular Nucleus
MiTg: Microcellular Tegmental Nucleus
MnR: Median Raphe
mRT: Midbrain Reticular Nucleus
MM: Mammillary Nucleus
MPL: Medial Paralemniscal Nucleus
PAG: Periaqueductal Gray
PMnR: Paramedian Raphe Nucleus
PnC: Pontine Reticular Nucleus, Caudal Part
PnO: Pontine Reticular Nucleus, Oral Part
PPTg: Pedunculopontine Tegmental Nucleus
PSth: Parasubthalamic Nucleus
PT: Pretectal Area
RI: Rostral Interstitial Nucleus of Medial Longitudinal Fasciculus
Rli: Rostral Linear Nucleus Raphe
RMg: Raphe Magnus
RPC: Red Nucleus, Parvocellular Part
RRF: Retrorubal Field
RtTg: Reticulotegmental Nucleus
SC: Superior Colliculus
SNc: Substantia Nigra Pars Compacta
SNl: Substantia Nigra Pars Lateralis
SNr: Substantia Nigra Pars Reticulata
SPTg: Subpeduncular Tegmental Nucleus
SubB: Sub–Brachial Nucleus
SuM: Supramammillary Nucleus
SuMx: Supramammillary decussation
VTA IF: Ventral Tegmental Area, Interfascicular Nucleus
VTA PBP: Ventral Tegmental Area, Parabrachial Pigmented Nucleus
VTA PN: Ventral Tegmental Area, Paranigral Nucleus
VTg: Ventral Tegmental Nucleus

